# Condition-specific series of metabolic sub-networks and its application for gene set enrichment analysis

**DOI:** 10.1101/200964

**Authors:** Van Du T. Tran, Sébastien Moretti, Alix T. Coste, Sara Amorim-Vaz, Dominique Sanglard, Marco Pagni

## Abstract

**Motivation:** Genome-scale metabolic networks and transcriptomic data represent complementary sources of knowledge about an organism’s metabolism, yet their integration to achieve biological insight remains challenging.

**Results:** We investigate here condition-specific series of metabolic sub-networks constructed by successively removing genes from a comprehensive network. The optimal order of gene removal is deduced from transcriptomic data. The sub-networks are evaluated via a fitness function, which estimates their degree of alteration. We then consider how a gene set, *i.e.* a group of genes contributing to a common biological function, is depleted in different series of sub-networks to detect the difference between experimental conditions. The method, named metaboGSE, is validated on public data for *Yarrowia lipolytica* and mouse. It is shown to produce GO terms of higher specificity compared to popular gene set enrichment methods like GSEA or topGO.

**Availability:** The *metaboGSE* R package is available at https://cran.r-project.org/web/packages/metaboGSE.

## 1 Introduction

The advent of high throughput sequencing techniques, especially RNA sequencing, has greatly facilitated the experimental investigation of an organism’s transcriptome under different physiological conditions. RNA-seq data consists of reads mapped onto an annotated genome, which permits quantitation of the transcript abundance of all predicted genes. These values can be used as a proxy to quantify gene expression or may provide hints about protein abundance for protein-coding genes. Differential expression between two conditions or co-expression profiles across many conditions are currently the fundamental statistical approaches to analyze RNA-seq data (Conesa *et al.*, 2016). However, to obtain a biological interpretation from these analyses, genes also need to be carefully annotated with prior biological knowledge. For example, the Gene Ontology (GO) provides genome annotations by grouping genes into sets, each one identified by a unique GO term that corresponds to a process, function or sub-cellular location. GO terms are hierarchically arranged in a directed acyclic graph (DAG) whose structure can be exploited by computational methods such as topGO (Alexa and Rahnenführer, 2016) together with gene expression data. Another popular method, GSEA (Subramanian *et al.*, 2005) considers two-condition expression profiles and attempts to identify functionally enriched sets of genes by investigating the change in the expression-based orderings of the genes. The incorporation of gene connectivity information has been shown as a way to improve gene set enrichment methods (Alexeyenko *et al.*, 2012; Glaab *et al.*, 2012). Such connectivity information could be provided by a genome scale metabolic network (GSMN).

GSMNs have been successfully used to study and model metabolism in living organisms (Feist and Palsson, 2008; McCloskey *et al.*, 2013; Oberhardt *et al.*, 2009). These are complex biological networks that include thousands of interconnected nodes of four main types: metabolites; biochemical and transport reactions; enzymes and transporters; and genes. The last three elements are usually referred to as GPR (gene-protein-reaction). A GSMN can be turned into a predictive tool to model if and how an organism can reach a certain objective, for instance biomass production, under defined environmental constraints. Flux Balance Analysis (FBA) (Varma and Palsson, 1994) can then be used for this purpose. For the sake of simplicity, we will designate here a model as *viable*, if it can produce biomass at a non-zero rate. A viable GSMN can be used to predict the essentiality of genes, for example, by simulating the effect on viability of a gene knockout, which can be further validated against experimental data (Imam *et al.*, 2015; O’Brien *et al.*, 2015; Simeonidis and Price, 2015). Model viability is sometimes referred to as model *consistency* in the literature.

Existing methods for integrating RNA-seq data with GSMN can be classified in two broad categories: those constraining FBA flux distribution in the network and those extracting context-specific sub-networks (Machado and Herrgård, 2014; Kim and Lun, 2014; Vivek-Ananth and Samal, 2016; Vijayakumar *et al.*, 2017; Opdam *et al.*, 2017). In Machado and Herrgård (2014), the first category is benchmarked and the authors concluded with two ambiguous observations: absolute and relative gene expression seem to perform similarly; and using gene expression as additional FBA constraints is not clearly superior to what is obtained with *parsimonious FBA*, an algorithm that does not depend on transcriptomic data at all. The second category, reviewed by Opdam *et al.* (2017), contains methods that: remove genes whose products are supposed to be absent in a given condition, by minimizing the flux through reactions with low gene expression (GIMME (Becker and Palsson, 2008)); optimize the trade-off between removing reactions with low gene expression and keeping reactions with high gene expression (iMAT (Zur *et al.*, 2010), INIT (Agren *et al.*, 2012)); keep an active set of core reactions while removing other reactions if possible (MBA (Jerby *et al.*, 2010), FASTCORE (Vlassis *et al.*, 2014), mCADRE (Wang *et al.*, 2012)). With all these methods, the choice of one or several cut-offs for gene expression strongly impact the submodel produced and its properties. Relatively little biological insight has been published from contrasting such context-specific sub-networks, with the exception of the predictions and validations of the flux on a few metabolites of interest (Machado and Herrgård, 2014) or gene essentiality (Opdam *et al.*, 2017).

The metaboGSE method presented here considers a whole series of sub-networks built by successively removing genes from an initial comprehensive network, rather than a single, possibly optimal sub-network to avoid the choice of a particular cut-off for gene expression. The optimal ranking of the genes to remove is determined as the most significant one when compared to random rankings, using the gene expression data. The viability of any sub-network is ensured by the introduction of artificial reactions and the minimization of the flux on them. This *rescue* procedure also allows for the formulation of a *fitness* function that measures how close an unviable sub-network is from a viable one. We then study how a gene set, *i.e.* a collection of genes contributing to a common biological function, is depleted in the series of sub-networks. Depletion curves are integrated with respect to fitness and tested for statistical significance given variations among experimental conditions and replicates.

To validate the method, we first used public experimental data and a metabolic model for *Yarrowia lipolytica*, a yeast which is widely exploited in industrial microbiology for lipid production (Ledesma-Amaro and Nicaud, 2016). The metabolic model iMK735 of *Y. lipolytica* (Kavšček *et al.*, 2015) was used, as it can simulate growth and production of lipids when oxygen consumption is limiting. Gene expression data were taken from the study of Maguire *et al.* (2014) on the role of the sterol-regulatory element binding protein Sre1 and the transcription factor Upc2 in sterol metabolism in hypoxic and normoxic conditions. A mouse dataset was also investigated, comprising RNA-seq data from macrophages in adipose tissue (Hill *et al.*, 2018) and the metabolic model iMM1415 (Sigurdsson *et al.*, 2010).

## 2 Materials and Methods

### 2.1 Datasets

The *Y. lipolytica* data analyzed is included in the *metaboGSE* R package from CRAN along with a vignette of the analysis pipeline. The mouse dataset is described in Supplementary Note S1.

#### 2.1.1 RNA-seq data

For *Y. lipolytica*, 22 RNA-seq samples of normoxic and hypoxic growth with *sre1*Δ*, upc2*Δ, *sre1*Δ */upc2*Δ mutants and the wild type strain were obtained from Maguire *et al.* (2014) (NCBI: PRJNA205557). These data are summarized in Table 1 and processed with standard preliminary RNA-seq data analysis (see Supplementary Note S3).

**Table 1.**
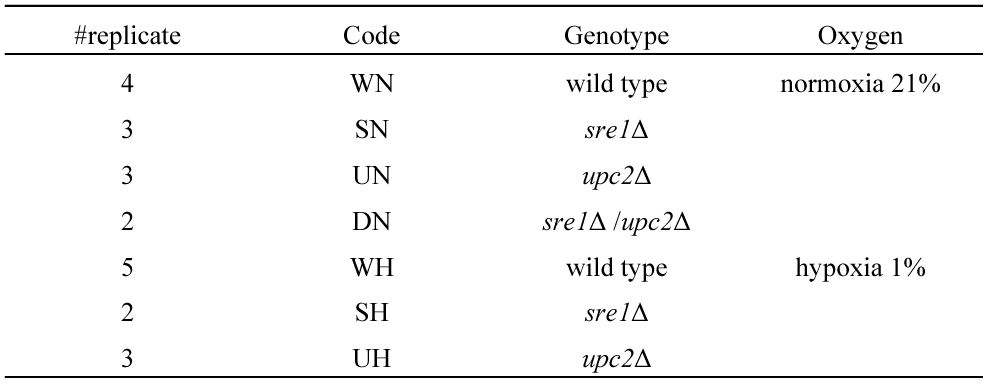
Designation of RNA-seq data obtained from Maguire *et al.*

#### 2.1.2 Genome-scale metabolic networks

The *Y. lipolytica* iMK735 model (http://www.ebi.ac.uk/biomodels, Kavšček *et al.*, 2015) with a production of 40% lipid content in the biomass was studied. The genome, proteome, GO annotations, and model were integrated within the framework of MetaNetX (Moretti *et al.*, 2016). The external reactions of the model were adapted to approximately simulate growth in Yeast extract Peptone Dextrose (YPD) medium in hypoxic (anaerobic) or normoxic (aerobic) environments (Maguire *et al.*, 2014) by modifying the oxygen supply. For normoxia, oxygen consumption was unrestricted as in the original model and took a value of 244 mmol•gDW^−1^•h^−1^ as given by the Minimum Total Flux algorithm where the sum of absolute values of fluxes was minimized. We then arbitrarily limited the available oxygen to 50 mmol•gDW^−1^•h^−1^ to simulate hypoxic conditions. Preliminary investigation showed that the model behavior did not significantly depend on the exact value of this setting. The model was cleaned by removing dead-end metabolites, which were only either produced or consumed. *Blocked* reactions as given by flux variability analysis (FVA) (Mahadevan and Schilling, 2003) were also removed, as explained below. The final investigated model contained 818 reactions, 469 genes, and 605 metabolites after cleaning, and was referred to as the *comprehensive* model in our analysis.

### 2.2 Metabolic sub-model construction

*Sub-model construction* by removing genes refers to the process of (i) removing an initial set of genes (input) and their associated reactions, (ii) determining the blocked reactions (see Supplementary Note S2), (iii) removing the blocked reactions with their associated genes, if any. A similar procedure named *deleteModelGenes* is available from the COBRA Toolbox (Heirendt *et al.*, 2017). Below, we refer to a particular sub-model with the number of initially removed genes, but all presented analysis results have been obtained after gene removal propagation through the blocked reactions.

#### 2.2.1 Measure of metabolic model fitness

Viability, for instance growth capacity, is crucial for the usability of a metabolic model. Removing a few reactions from a viable network is often sufficient to render it unviable. Here we propose a measure to assess how close an unviable model is to a viable one, which will be defined as the *fitness* of the model.

##### 2.2.1.1 Principle of growth rescue

In this section, we introduce a procedure to restore the viability of an unviable metabolic network by the introduction of artificial reactions and minimizing the flux on them. It consists first in modifying the input network around the growth reaction as illustrated in Figure 1. An artificial metabolite is created to replace each of those present in the growth reaction except biomass itself. Each artificial metabolite *x’* is linked to the original metabolite *x* through a directed *help* reaction denoted *h_x_*. In addition, *x’* can be produced or consumed via a *rescue* external reaction denoted *r_x_*. Note that the purpose of any help reaction is to avoid artificially supplying *x* as a side effect of rescuing *x’*. No constraints are applied on the fluxes of the rescue and help reactions apart from their directions. The same network modifications could be applied to the non-growth associated maintenance reaction that does not allow for a zero flux.

**Fig. 1.**
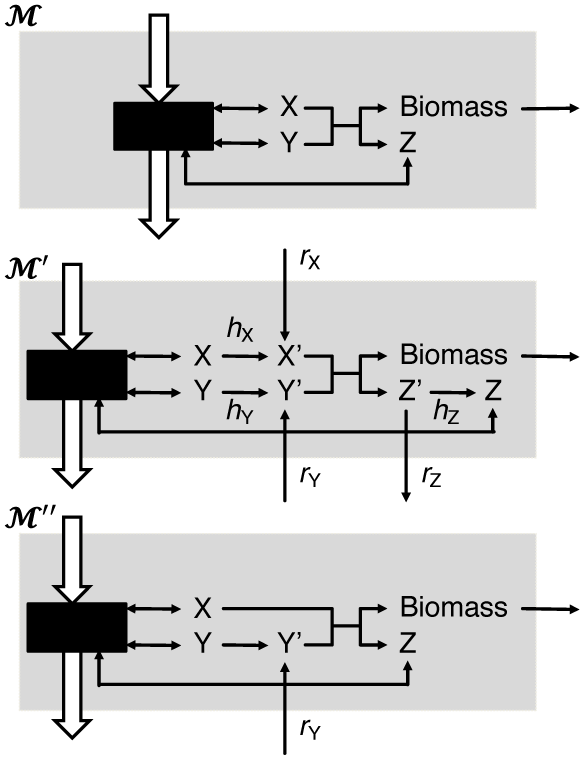
Schema of GSMN rescue process. 𝓜, original GSMN with growth reaction X + Y ---> Z + Biomass. *𝓜*′, expanded GSMN with the full set of rescue (*r_x_*) and help (*h_x_*) reactions for every metabolite x in the biomass reaction. *𝓜*″, example of a minimal rescued GSMN in the particular case where only metabolite Y needs to be rescued.

Let *𝓜′* be the expanded version of a GSMN *𝓜* with the modified growth reaction and the full set of rescue and help reactions, **S′** is the corresponding stoichiometric matrix (see basic notions on GSMN in Supplementary Note S2). The flux distribution ***𝑣′*** is constrained by the bound vectors **Ib′** and **ub′** that account for the directionality of newly added reactions and constraint the growth reaction at a fixed rate, arbitrarily chosen as 20% the original model growth objective, *i.e.* 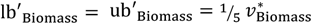.

Let **c**_Rescue_ be a vector of coefficients that are equal to 0 for all but the rescue reactions that receive a coefficient of 1/*B*_*x*_ where *B_x_* is the stoichiometric coefficient for the metabolite *x* in the original growth reaction. The rescue procedure can be stated as the following linear programming (LP) problem:

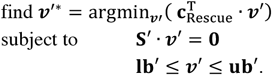

Only rescue reactions for metabolite *x* with non-zero flux 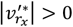 are required to restore the model viability. A viable model *𝓜″* with a minimal flux of rescue reactions could be formulated as shown in Figure 1. It must be noted that the above LP problem could admit more than one solution ***𝑣′\**** and different solutions might be expected if **c**_Rescue_ was set differently. We investigated this problem through simulation with the iMK735 model and observed that most of the solutions are unique and do not depend on the precise values of **c**_Rescue_.

Figure 2A presents the fraction of metabolites in the growth reaction that need to be rescued after randomly removing genes in the model. This simulation shows the crucial property that the more genes removed, the more the model is damaged. Figure 2B presents the fraction of random models where each individual metabolite needs to be rescued, clearly showing different behaviors among metabolites in the growth reaction. This suggests that the different metabolites should not be treated equally.

**Fig. 2.**
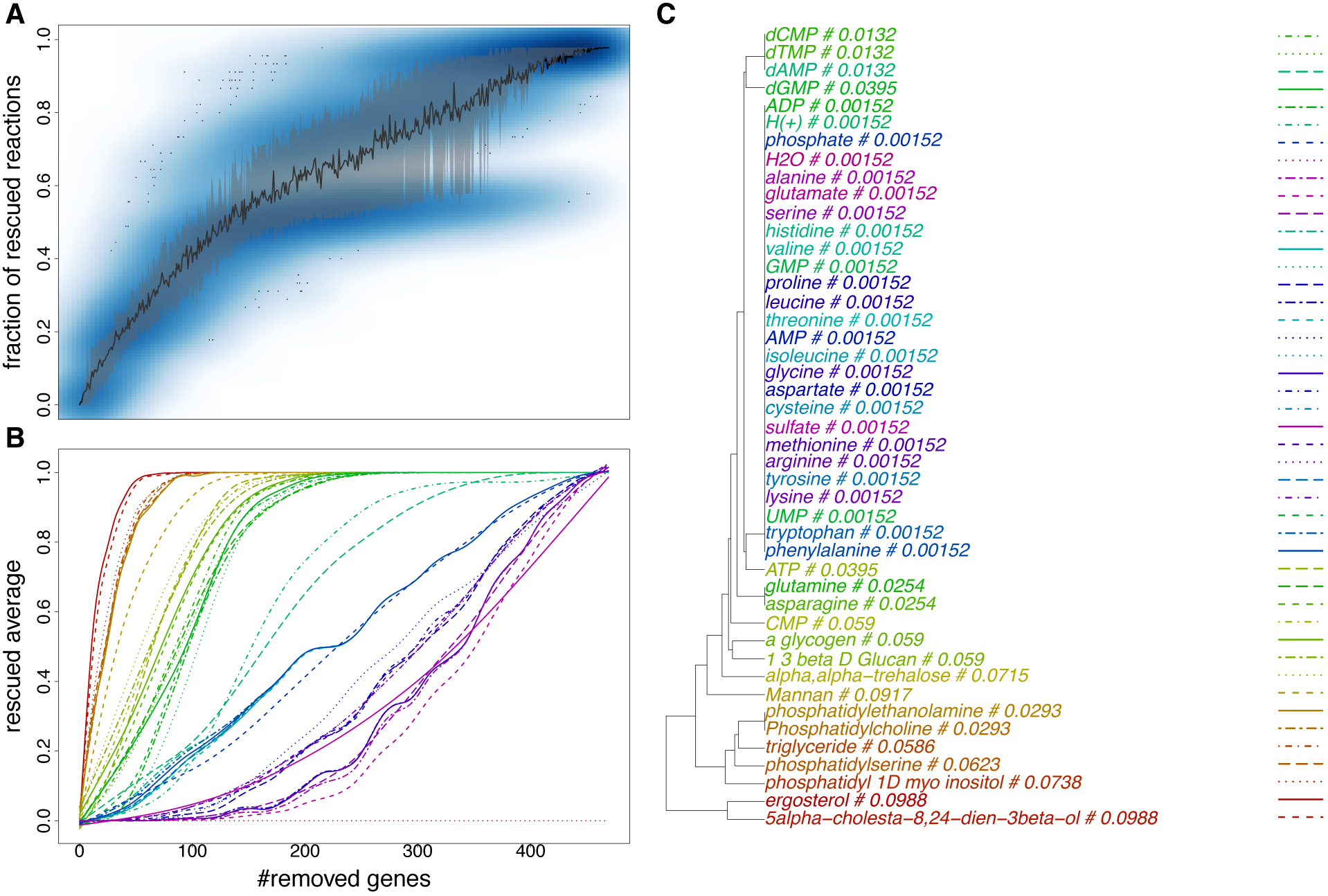
Rescue in metabolic sub-models obtained by removing different numbers of random genes (N = 50 draws) from *Y. lipolytica* model iMK735. (A) Fraction of metabolites that need to be rescued via corresponding rescue reactions. Blue darkness in the scatter plot represents the density of fractions of rescued reactions. Black curve indicates average fractions. Gray region represents 20%-80% quantiles of random fractions. (B) Fraction of draws where each individual metabolite is rescued. (C) Weights of rescue reactions obtained via our weighting schema.

##### 2.2.1.2 Weighting scheme for model fitness

A weighting scheme is introduced to account for the variable importance of metabolites in the growth reaction and possible dependencies among them. Model fitness is defined as the realized objective of the following LP problem:

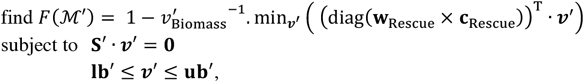

Where 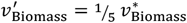, diag(A) is the diagonal of a square matrix A, **w**_Rescue_ is a vector of weights that are equal to 0 for all but the rescue reactions and that are normalized to sum to 1. **w**_Rescue_ is computed in the following procedure. For every gene in the model, a single gene knockout is simulated, and the rescue procedure is performed to determine which metabolites in the growth reaction are affected, using the previously presented LP problem. Hence, every rescue reaction *r_x_* can be associated with a binary vector which describes whether the reaction is needed to rescue each of the gene knockouts. These binary vectors are used to compute Euclidean distances between rescue reactions, which are used in hierarchical clustering with average linkage, and the Gerstein method (Gerstein *et al.*, 1994) is then applied to the resulting tree as a means to assign a weight to each rescue reaction. Those rescue reactions with similar binary vectors share weights, while those with unique profile receive a larger weight. The weighting schema has two main effects: (i) it reduces the importance of metabolites that are hardly affected by slightly damaging the model, and (ii) assigns similar importance to metabolites that appear on the same pathway, as shown in Figure 2C. For instance, H_2_O has the smallest weight, ergosterol and 5alpha-cholesta-8,24-dien-3beta-ol in the same pathway share the same weight. Other sampling schemes were investigated, for example by removing several genes at once, but these did not yield very different metabolite weights. Hence the simpler experimental setting was used. The simulation of random gene removal in Figure 2A and 3 shows that the introduction of this weighting scheme produced the fitness scores that are less dispersed than the fraction of rescued reactions.

#### 2.2.2 Optimal ranking of genes for removal

The proposed fitness function can be used to evaluate a series of condition-specific sub-models constructed by removing genes in any order. The question then arises as to how to optimally rank genes for removal, such as minimizing network disruption *i.e.* preserving its fitness. In this section, we investigate different metrics to rank genes according to their expression in a given experiment.

We tested the following transformations of the raw expression data to rank the genes:

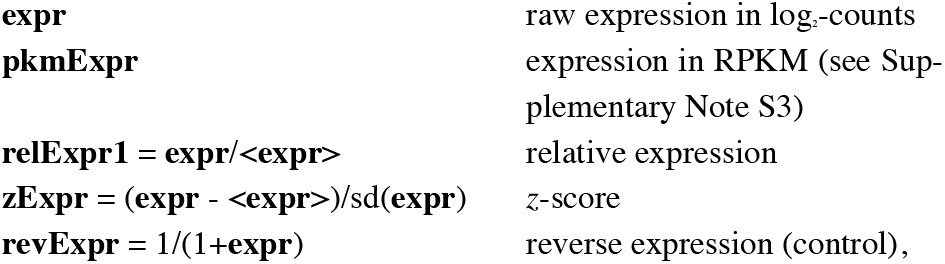

where <**expr**> and sd(**expr**) denote the average and standard deviation across conditions, respectively.

Let *ρ* be a ranking of the genes. Sub-models are constructed by removing genes in the order given by *ρ*. The resulting fitness scores decrease when more genes are removed, and the decreasing trend depends on *ρ*. Figure 3 illustrates such reductions obtained by successively removing genes from the comprehensive model iMK735 for the UH condition (*upc2*Δ in hypoxic condition, UH2 sample). In this example **expr** is the most fitness-preserving ranking and **revExpr** is the worst one.

**Fig. 3.**
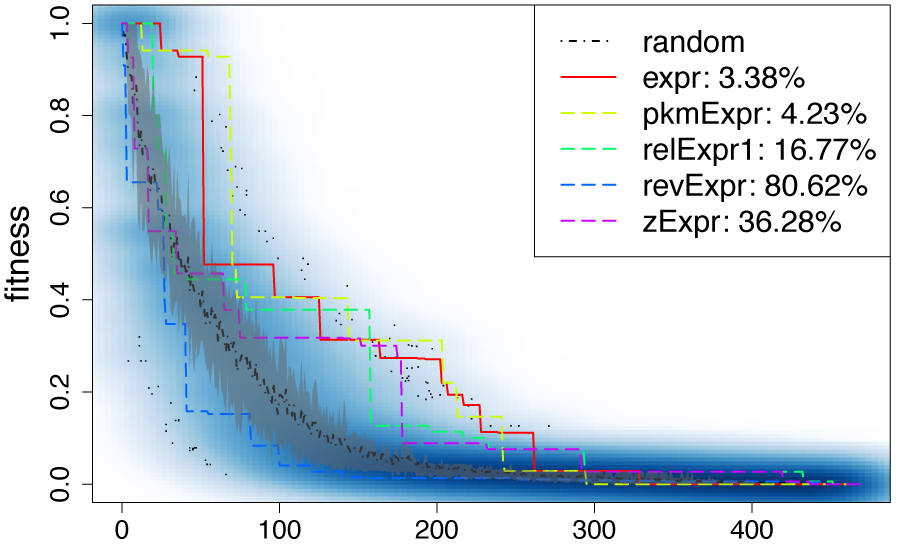
Fitness decrease while removing genes from the iMK735 model with different rankings in the UH condition (UH2 sample). **random**: random draw, **expr**: voom-normalized expression, **pkmExpr**: voom-normalized expression in RPKM, **relExpr1**: relative expression **expr/<expr>**, **revExpr**: reverse expression, **zExpr**: *z*-score. Blue darkness in the scatter plot represents the density of random fitness. Gray region represents 20%-80% quantiles of random fitness.

We assess a ranking *ρ* by *performance index Z_ρ_*, which indicates the percentage of random draws yielding sub-model fitness higher than that of sub-models created by *ρ*-based removal of the same number of genes and which is weighted by the average random fitness. *Z_ρ_* is computed as follows:

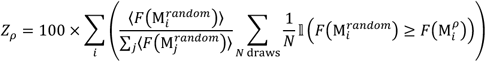

where 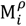 denotes the rescued sub-model of the comprehensive network M obtained after removing the *i* genes using ranking *ρ* and 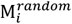 the rescued sub-model by randomly removing *i* genes, 〈*F*(…)〉 the average fitness on *N* draws. The lower the performance index, the better the ranking. The optimal ranking is the one dominating the others by producing the lowest performance index for all investigated RNA-seq samples. For instance, the absolute expression ranking **expr** is determined as the best one for 19 of 22 the *Y. lipolytica* samples, whereas it is **pkmExpr** for the mouse dataset (see Table S1). Other rankings such as (**expr**^2^/<**expr**>)^1/2^ and (**expr**^3^/<**expr**>)^1/3^ were also investigated, yet not comparable to the selected ones. In the rest of this article, the **expr** and **pkmExpr** rankings will be used in the *Y. lipolytica* and mouse study, respectively.

### 2.3 metaboGSE: contrasting gene set enrichment in condition-specific sub-models

We introduce here the metaboGSE method, which aims at identifying gene sets that are differentially enriched. The method consists of three steps depicted in Figure 4 and illustrated in Figure 5 for GO:0006635 – fatty acid beta-oxidation. The first step consists in constructing a series of sub-models for every sample and computing their fitness profile.

**Fig. 4.**
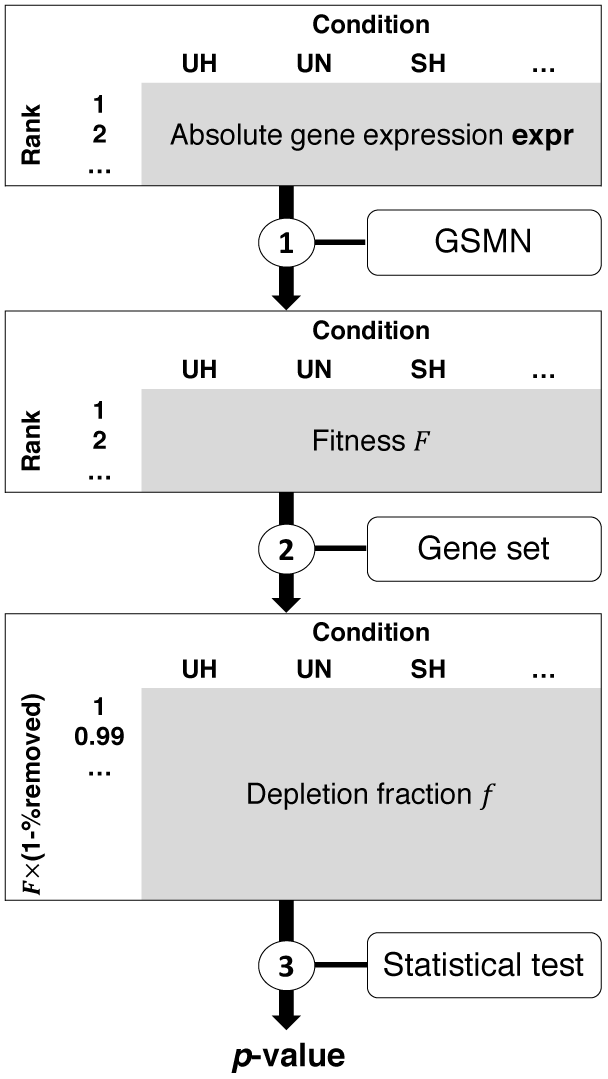
metaboGSE algorithm.

**Fig. 5.**
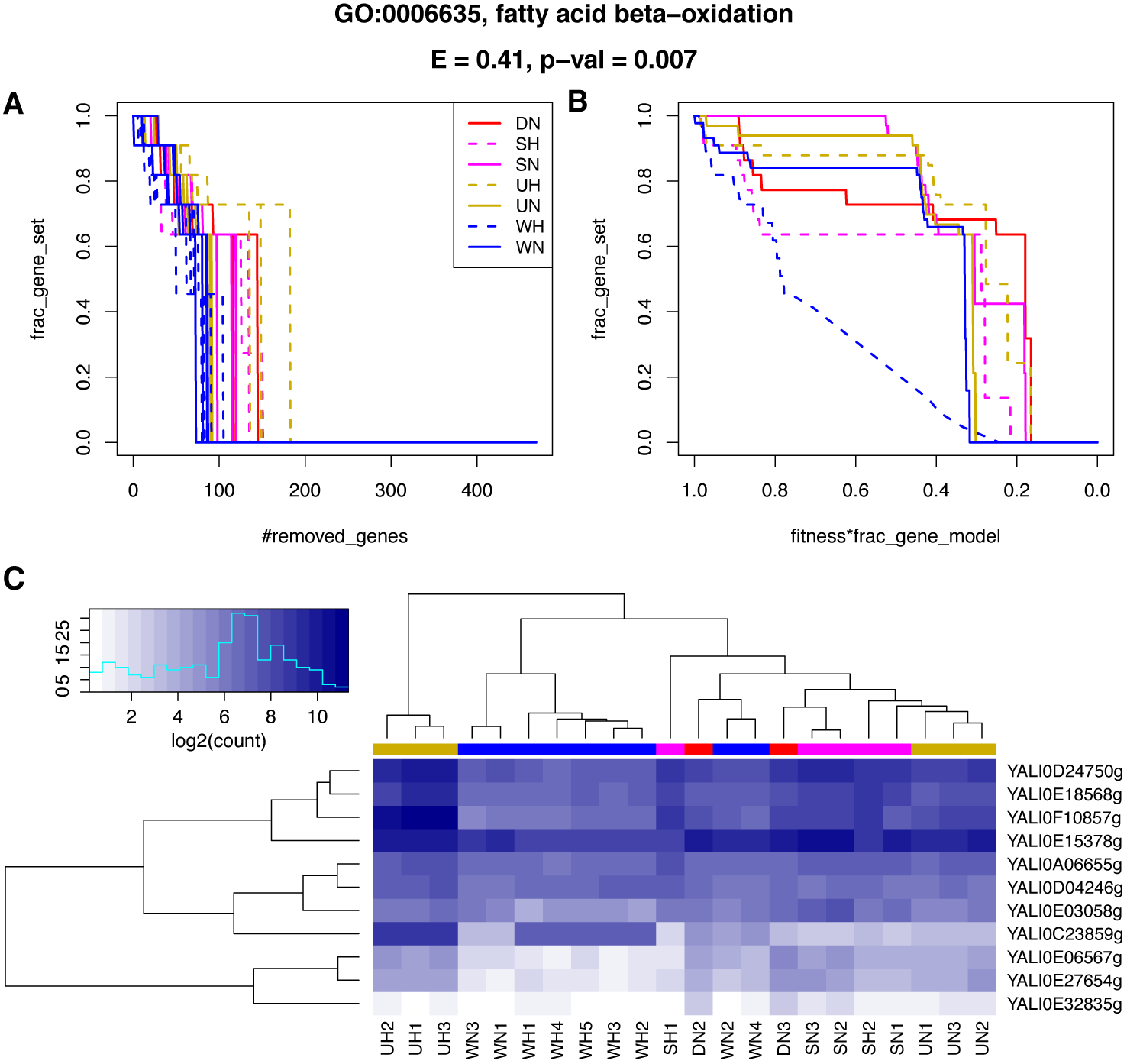
Enrichment of GO:0006635 fatty acid beta-oxidation in *Y. lipolytica* sub-models in seven RNA-seq conditions. (A) Depletion fraction in function of number of removed genes (B) Depletion curve: depletion fraction in function of fitness times fraction of remaining genes in the model (C) Expression of associated genes

In the second step, for a given gene set *g*, we compute the *depletion fraction* 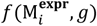, *i.e.* fraction of *g*-associated genes remaining in each sub-model, where 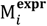 denotes the rescued sub-model after the removal of *i* genes. The evolution of 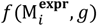 is plotted as a function of 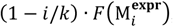, defining a depletion curve, where *k* denotes the total number of genes in M. The depletion curve for each condition is the average curve on all replicates. As shown in Figure 5A, the fraction *f* of genes associated with fatty acid beta-oxidation decreases rapidly in all conditions, but the depletion curve clearly separates hypoxic wild-type from the other conditions (Figure 5B). The down-regulation of several but not all GO:0006635 genes in the WH condition is illustrated in Figure 5C.

In the third step, we perform a permutation test for the significance of discrepancy of the given gene set *g* across conditions. The test statistic is defined as the maximum area between every pair of depletion curves among all conditions. The resampling is performed by permuting replicates between conditions while keeping unchanged the number of replicates in each condition. The resulting *p*-value indicates whether the depletion evolution of *g* in one condition differs from at least one of the others. For GO:0006635, the discrepancy of WH versus the other conditions is justified with a *p*-value of 0.007 and a test statistic of 0.41 (Figure 5). These *p*-values are subsequently adjusted by Benjamini-Hochberg (BH) correction across all the studied gene sets (Benjamini and Hochberg, 1995). A similar post-hoc permutation test is also implemented for pairwise comparisons between conditions to check for the pairwise differential signal.

## 3 Results

### 3.1 Condition-specific sub-model construction on *Y. lipolytica* data

We applied the selected gene removal strategy to construct sub-models for the seven conditions in Maguire *et al.* (2014) (Table 1). The **expr** rankings are different between conditions despite a similar distribution of gene expression values (Figure 6A). The sub-model series constructed based on **expr**, and thus their fitness, evolve differently across conditions (Figure 6B). Figure 6C-F shows a high degree of variation across conditions in the number of genes and reactions (post propagations through blocked reactions), as well as in the fraction of genes and reactions that are essential. Besides, the erratic evolutions of essential genes and reactions is noteworthy and might be associated with the unexpectedly large spread in essentiality predictions recently reported in Opdam *et al.* (2017).

**Fig. 6.**
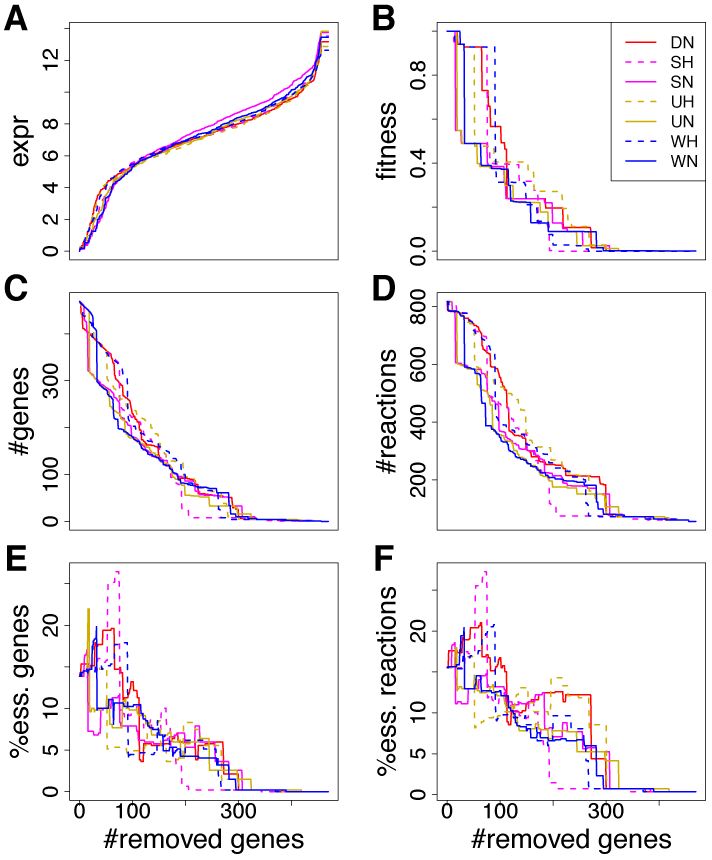
Gene expression (A), fitness (B), number of genes (C), number of reactions (D), percentage of essential genes (E), and percentage of essential reactions (F) of *Y. lipolytica* sub-models from seven RNA-seq conditions. obtained by removing genes following the **expr** ranking. Only one replicate from each condition is shown. The percentages represent the numbers of essential reactions or genes in sub-models over those in the comprehensive model.

To compare the condition-specific sub-model construction from the existing methods with that from our approach, we investigated GIMME (Becker and Palsson, 2008) and iMAT (Zur *et al.*, 2010). These two methods were benchmarked in Opdam *et al*. (2017) and could be used with information that is deduced only from GSMN and transcriptomics data. We built a sequence of sub-models with GIMME using 12 gene expression cut-offs (in log2 RPKM) from 0 to 11 for each of the 22 samples. For iMAT, these cut-offs were used as *threshold_lb* while *threshold_ub* was determined as *threshold_lb + 2*standard_deviation(expression)*, as recommended in the Supplementary Data of Zur *et al*. (2010). The choice of such a limited number of cut-offs was due to highly time-consuming model construction process of the two methods in Matlab. Indeed, the parallelization implemented in metaboGSE using the *sybil* and *parallel* R packages allowed us to construct the complete list of all sub-models in almost one hour on a 64-processor Intel Xeon E5-4620 of 2.6 GHz. The MatLab implementations of iMAT and GIMMME were much slower in our hands. Intersections and unions of genes in each 22 sub-models were investigated to evaluate the difference between them. Figure S1 shows 1 – Intersection/Union plotted as a function of *k* – Union, where *k* denotes the number of genes in the comprehensive GSMN. Interestingly, the submodels produced with metaboGSE were quite similar to those produced by iMAT, a method that relies on a MILP algorithm. GIMME produced sub-models that were less distinct across conditions.

### 3.2 Gene set enrichment on *Y. lipolytica* data

To validate the biological findings produced by our approach, we investigated 135 gene sets defined as *GO135* (see Supplementary Note S4). Maguire *et al*. (2014) studied the role of Sre1 and Upc2 in regulating sterol metabolism in hypoxic and normoxic conditions in *Y. lipolytica* by performing GO term enrichment analysis of differentially expressed genes using DAVID. Among the 116 biological-process GO terms they reported, only eight were found in *GO135*, but none had a reported BH-adjusted *p*- value < 0.15. Here we compare our results with those of topGO *weight01* and GSEA on the iMK735 genes (see Supplementary Note S4).

*Condition contrast is predominantly hypoxia-normoxia*. The depletion curves of the top 50 GO terms found to be significantly enriched with metaboGSE (ordered by permutation test statistic with FDR < 0.05) (see Figure S2) suggest that the degree of oxygen limitation is the most likely explanation for the enrichment of 36 GO terms, which is expected for this dataset – where hypoxia is the only environmental variable. Interestingly, for 24 of them, normoxic double-mutant condition is likely grouped with hypoxic conditions while contrasting with other normoxic conditions.

*metaboGSE reveals GO terms of higher specificity*. A GO term is considered of higher specificity compared to another one when it is an offspring of the latter (Ashburner *et al.*, 2000). The top 50 GO terms found to be significantly enriched by metaboGSE (ordered by permutation test statistic with FDR < 0.05), topGO, and GSEA (ordered by FDR) unite into 99 GO terms and are summarized in Figure S3 and Table S2. These 99 terms also included the eight found in Maguire *et al*. All terms from metaboGSE are of higher or equal specificity, *i.e.* offspring or identical, to those found by topGO and/or GSEA. Thirty-five among the 50 GO terms from metaboGSE are related to those found by the other methods yet include 9 terms of higher specificity: GO:0034637 (cellular carbohydrate biosynthetic process), GO:0015937 (coenzyme A biosynthetic process), GO:0071265 (L-methionine biosynthetic process), GO:0046474 (glycerophospholipid biosynthetic process), GO:0097164 (ammonium ion biosynthetic process), GO:0006656 (phosphatidyl choline biosynthetic process), GO:0001676 (long-chain fatty acid metabolic process), GO:0043649 (dicarboxylic acid catabolic process), and GO:0009098 (leucine biosynthetic process). Six of them are found in the three largest connected DAGs of enriched GO terms depicted in Figure 7.

**Fig. 7.**
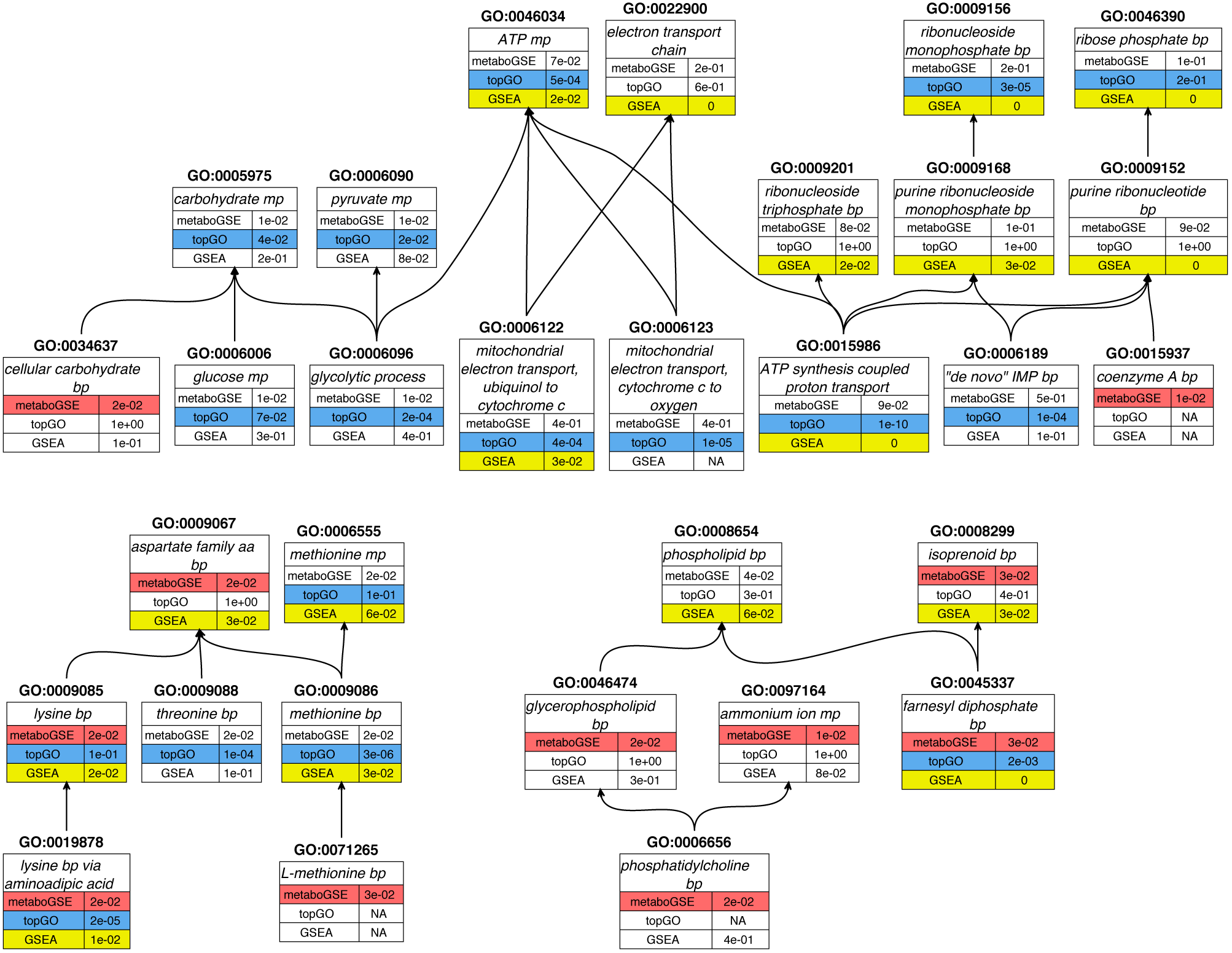
The three largest connected DAGs enriched by either metaboGSE (in red), topGO weight01 (in blue) or GSEA (in yellow). Table S2 provides the full listing of the 99 GO terms resulting from the union of the top 50 GO terms from each method. FDR is reported for each GO term. NA value indicates that the GO term was not reported by the corresponding method. bp: biosynthetic process, cp: catabolic process, mp: metabolic process.

*Difference in sterol biosynthesis is confirmed.* Ergosterol biosynthetic process (GO:0006696) is enriched by all methods. Along with GSEA and topGO, we also discover GO:0045337 (farnesyl diphosphate biosynthetic process) (Figure 7), which is part of the ergosterol synthesis pathway. This reflects the experimental design investigated in Maguire *et al*. (2014), *i.e.* the regulation of sterol metabolism. In Figure S4AB, the sudden drop from 100% to 0% of 19 ergosterol-associated genes indicates that either all these genes or none of them exist in each sub-model. This sudden drop is caused by the propagation through the blocked reactions in the linear pathway of ergosterol biosynthesis (Parks and Casey, 1995). Interestingly, a detailed manual investigation of this case revealed that the sudden drops in the different conditions are actually caused by the altered ranks of either YALI0D17050g, YALI0E18634g or YALI0F26323g, which do not belong to GO:0006696 gene set. Figure S4C presents the expression profiles of all those genes.

*metaboGSE identifies contrasts other than hypoxia-normoxia*. All GO terms relating to fatty acid metabolism (GO:0001676 long-chain fatty acid metabolic process, GO:0009062 fatty acid catabolic process, GO:0006635 fatty acid beta-oxidation, GO:0006631 fatty acid metabolic process) show an earlier drop in their depletion curve in the hypoxic wildtype comparing to the remaining conditions that stay similar (Figure S2). This might suggest that the presence of both Sre1 and Upc2 in low-oxygen growth condition reduces the expression of YALI0E03058g, which causes the earlier drop in WH (Figure 5).

### 3.3 Application of metaboGSE to mouse data

metaboGSE was also applied to the mouse dataset from Hill *et al.* (2018) using the iMM1415 model (Sigurdsson *et al.*, 2010) (see Supplementary Note S1). The **pkmExpr** ranking was figured as the best ranking among those simulated (see Table S1). Only four genes of iMM1415 were found to be differentially expressed between PBS and Ly6C (fold-change ≥ 2, FDR < 0.05), resulting in no significantly enriched GO terms with topGO *weight01* and GSEA. We then applied these two methods to all differentially expressed genes in the genome, but not only to those in iMM1415, to increase the number of differentially expressed genes. We scrutinized a list of 24 GO terms related to inflammatory response, cholesterol and lipid biosynthesis as reported in Hill *et al.* (2018) and associated to at least one iMM1415 gene. The results shown in Figure S5 reveal that metaboGSE can detect GO terms that are located lower in the Gene Ontology, like GO:0050728 (negative regulation of inflammatory response), GO:0002675 (positive regulation of acute inflammatory response), and GO:1903725 (regulation of phospholipid metabolic process), despite working on a much smaller collection of genes than the other two methods. This confirms the ability of metaboGSE to capture GO terms of higher specificity as observed above with *Y. lipolytica*.

## 4 Discussion

We present here a method for gene set enrichment analysis that utilizes a GSMN as an additional source of information and that focuses on genes expressed at low level. Our central working hypothesis is that the correlation between gene expression levels and fluxes on related reactions is very poor in general, but the low expressed genes are plausibly associated with zeroed fluxes. This method is complementary to established methods such as topGO and GSEA that focus on differential expression of sufficiently expressed genes. The introduction of a GSMN restricts the list of investigated genes to those present in the model (*i.e.* related to metabolism), and thus the list of gene sets that can be discovered. The formulation of the external reactions of the GSMN and how well they represent the experimental system are likely to affect metaboGSE outcome, although this has not been investigated in detail here. Tissue-specific models are not a prerequisite to utilize metaboGSE. The GSMN and the set of RNA-seq data both need to be of high quality and adequate for experimental designs. Our method is capable of producing more informative GO terms (*i.e.* that are located lower in the Gene Ontology) than those returned by GSEA and topGO for example. This might be because metaboGSE can increase the size of investigated gene sets by considering structural constraints brought by the propagation through blocked reactions, as for example the linearity of the ergosterol biosynthesis pathway in the case reported here. The genes affecting the discrepancy between conditions, which are not necessarily differentially expressed, can be further investigated for each enriched gene set. metaboGSE produces biologically meaningful results to the extent one can interpret them.

Our method does not aim at producing a condition-specific sub-model, but rather integrates on a series of them, thus avoiding the choice of a particular number of genes to remove. A GSMN is a drastic simplification of our understanding and knowledge of biochemistry that neglects most kinetic aspects in its representation of metabolism. On the modeling level, defining a sub-model by removing genes is equivalent to a gene knockout obtained from a molecular construct. It is likely that the metabolism dominating a given physiological state owes more to kinetic regulation than can be accounted for by only the metabolism structure. Moreover, it is very hard to ascertain that a gene is not expressed at all and even in this case, the absence of mRNA does not exclude that the protein is still present at a low concentration, as a remnant of a previous growth phase. Likewise, the presence of the protein does not ensure it is active. The construction of a series of sub-models followed by their rescue is essentially a way to circumvent the hard constraint caused by model viability and exploit sub-model properties that would be out of range. Our method to construct metabolic sub-networks could also be performed with other omics data, such as proteomics and metabolomics, and applied to other research problems.

The fitness function is the key component of our method. The proposed measure of fitness shows its capacity to capture the health status of a submodel and thus suggests some control on our sub-model construction. Despite the biologically meaningful results obtained with the datasets studied, several lines of improvement can be envisaged in future work, including: the formulation of the growth reaction could be improved by considering more metabolites; the proposed weighting scheme is likely suboptimal; and other dynamic properties of the model could be considered separately from the score derived from the LP-based minimization.

## Acknowledgements

We thank Ioannis Xenarios, Mark Ibberson, Frédéric Burdet, Alan Bridge, Nicolas Guex, Brian Stevenson, Frédéric Schütz, Mathias Ganter, and Jörg Stelling for helpful discussions and supports at different stages of the project, Olivier Martin for supports in MetaNetX development, and Claus Jonathan Fritzemeier for supports with *sybil*.

## Funding

This work was supported by the Swiss National Science Foundation grant [CRSII3_141848] (Sinergia) to Dominique Sanglard, Salomé Leibundgut and Marco Pagni. This work was also supported in part by the SystemsX.ch initiative under the projects MetaNetX and HostPathX. The computations were performed at the Vital-IT Center for high-performance computing (https://www.vital-it.ch) of the SIB Swiss Institute of Bioinformatics. SIB receives financial supports from the Swiss Federal Government through the State Secretariat for Education and Research (SER).

## Conflict of Interest

none declared.

## References

Agren, R. et al. (2012) Reconstruction of genome-scale active metabolic networks for 69 human cell types and 16 cancer types using INIT. PLoS Comput. Biol., 8, e1002518.

Alexa, A. and Rahnenführer, J. (2016) topGO: Enrichment Analysis for Gene Ontology. R package version 2.24.0.

Alexeyenko, A. et al. (2012) Network enrichment analysis: extension of gene-set enrichment analysis to gene networks. BMC Bioinformatics, 13, 226.

Ashburner, M. et al. (2000) Gene Ontology: tool for the unification of biology. Nat. Genet., 25, 25–29.

Becker, S.A. and Palsson, B.O. (2008) Context-specific metabolic networks are consistent with experiments. PLoS Comput. Biol., 4, e1000082.

Benjamini, Y. and Hochberg, Y. (1995) Controlling the False Discovery Rate: A Practical and Powerful Approach to Multiple Testing. J. R. Stat. Soc. Ser. B Methodol., 57, 289–300.

Conesa, A. et al. (2016) A survey of best practices for RNA-seq data analysis. Genome Biol., 17.

Feist, A.M. and Palsson, B.Ø. (2008) The growing scope of applications of genome-scale metabolic reconstructions using Escherichia coli. Nat. Biotechnol., 26, 659–667.

Gerstein, M. et al. (1994) Volume changes in protein evolution. J. Mol. Biol., 236, 1067–1078.

Glaab, E. et al. (2012) EnrichNet: network-based gene set enrichment analysis. Bioinformatics, 28, i451–i457.

Heirendt, L. et al. (2017) Creation and analysis of biochemical constraint-based models: the COBRA Toolbox v3.0. ArXiv171004038 Q-Bio.

Hill, D.A. et al. (2018) Distinct macrophage populations direct inflammatory versus physiological changes in adipose tissue. Proc. Natl. Acad. Sci. U. S. A., 115, E5096–E5105.

Imam, S. et al. (2015) Data-driven integration of genome-scale regulatory and metabolic network models. Front. Microbiol., 6, 409.

Jerby, L. et al. (2010) Computational reconstruction of tissue-specific metabolic models: application to human liver metabolism. Mol. Syst. Biol., 6, 401.

Kavšček, M. et al. (2015) Optimization of lipid production with a genome-scale model of Yarrowia lipolytica. BMC Syst. Biol., 9, 72.

Kim, M.K. and Lun, D.S. (2014) Methods for integration of transcriptomic data in genome-scale metabolic models. Comput. Struct. Biotechnol. J., 11, 59–65.

Ledesma-Amaro, R. and Nicaud, J.-M. (2016) Yarrowia lipolytica as a biotechnological chassis to produce usual and unusual fatty acids. Prog. Lipid Res., 61, 40–50.

Machado, D. and Herrgård, M. (2014) Systematic Evaluation of Methods for Integration of Transcriptomic Data into Constraint-Based Models of Metabolism. PLOS Comput Biol, 10, e1003580.

Maguire, S.L. et al. (2014) Zinc finger transcription factors displaced SREBP proteins as the major Sterol regulators during Saccharomycotina evolution. PLoS Genet., 10, e1004076.

Mahadevan, R. and Schilling, C.H. (2003) The effects of alternate optimal solutions in constraint-based genome-scale metabolic models. Metab. Eng., 5, 264–276.

McCloskey, D. et al. (2013) Basic and applied uses of genome-scale metabolic network reconstructions of Escherichia coli. Mol. Syst. Biol., 9, 661.

Moretti, S. et al. (2016) MetaNetX/MNXref - reconciliation of metabolites and biochemical reactions to bring together genome-scale metabolic networks. Nucleic Acids Res., 44, D523–526.

Oberhardt, M.A. et al. (2009) Applications of genome-scale metabolic reconstructions. Mol. Syst. Biol., 5, 320.

O’Brien, E.J. et al. (2015) Using Genome-scale Models to Predict Biological Capabilities. Cell, 161, 971–987.

Opdam, S. et al. (2017) A Systematic Evaluation of Methods for Tailoring GenomeScale Metabolic Models. Cell Syst., 4, 318–329.e6.

Parks, L.W. and Casey, W.M. (1995) Physiological implications of sterol biosynthesis in yeast. Annu. Rev. Microbiol., 49, 95–116.

Sigurdsson, M.I. et al. (2010) A detailed genome-wide reconstruction of mouse metabolism based on human Recon 1. BMC Syst. Biol., 4, 140–140.

Simeonidis, E. and Price, N.D. (2015) Genome-scale modeling for metabolic engineering. J. Ind. Microbiol. Biotechnol., 42, 327–338.

Subramanian, A. et al. (2005) Gene set enrichment analysis: a knowledge-based approach for interpreting genome-wide expression profiles. Proc. Natl. Acad. Sci. U. S. A., 102, 15545–15550.

Varma, A. and Palsson, B.O. (1994) Metabolic Flux Balancing: Basic Concepts, Scientific and Practical Use. Nat. Biotechnol., 12, 994–998.

Vijayakumar, S. et al. (2017) Seeing the wood for the trees: a forest of methods for optimization and omic-network integration in metabolic modelling. Brief. Bioinform.

Vivek-Ananth, R.P. and Samal, A. (2016) Advances in the integration of transcriptional regulatory information into genome-scale metabolic models. Biosystems, 147, 1–10.

Vlassis, N. et al. (2014) Fast Reconstruction of Compact Context-Specific Metabolic Network Models. PLoS Comput. Biol., 10.

Wang, Y. et al. (2012) Reconstruction of genome-scale metabolic models for 126 human tissues using mCADRE. BMC Syst. Biol., 6, 153.

Zur, H. et al. (2010) iMAT: an integrative metabolic analysis tool. Bioinforma. Oxf. Engl., 26, 3140–3142.

